# Nanopore sequencing from protozoa to phages: decoding biological information on a string of biochemical molecules into human-readable signals

**DOI:** 10.1101/2024.08.04.606558

**Authors:** Branden Hunter, Timothy Cromwell, Hyunjin Shim

## Abstract

Biological information is encoded in a sequence of biochemical molecules such as nucleic acids and amino acids, and nanopore sequencing is a long-read sequencing technology capable of directly decoding these molecules into human-readable signals. The long reads from nanopore sequencing offer the advantage of obtaining contiguous information, which is particularly beneficial for decoding complex or repetitive regions in a genome. In this study, we investigated the efficacy of nanopore sequencing in decoding biological information from distinctive genomes in metagenomic samples, which pose significant challenges for traditional short-read sequencing technologies. Specifically, we sequenced blood and fecal samples from mice infected with *Trypanosoma brucei*, a unicellular protozoan known for its hypervariable and dynamic regions that help it evade host immunity. Such characteristics are also prevalent in other host-dependent parasites, such as bacteriophages. The taxonomic classification results showed a high proportion of nanopore reads identified as *T. brucei* in the infected blood samples, with no significant identification in the control blood samples and fecal samples. Furthermore, metagenomic de novo assembly of these nanopore reads yielded contigs that mapped to the reference genome of *T. brucei* in the infected blood samples with over 96% accuracy. This exploratory work demonstrates the potential of nanopore sequencing for the challenging task of classifying and assembling hypervariable and dynamic genomes from metagenomic samples.

## Introduction

Decoding heritable biological information from a string of biochemical molecules, such as DNA, into human-readable signals involves several key steps in molecular biology and bioinformatics [1,2]. Until recently, short-read sequencing technologies utilize massively parallel sequencing to rapidly and accurately decode large volumes of DNA by generating millions of short-read sequences simultaneously [3]. However, these technologies require an amplification step, such as PCR, to produce the large quantities of DNA needed for massively parallel sequencing; this amplification is necessary for signal detection but prevents the direct sequencing of native DNA [4]. Practically, this amplification introduces potential biases, errors, and loss of important epigenetic modifications inherent in the original DNA sequence, while also preventing the direct conversion of biological information to human-readable signals.

Nanopore sequencing technology allows for the direct conversion of biochemical signals into human-readable sequences by exploiting the biophysical properties of nanopores and leveraging advanced computational methods (Figure 1A). As a single DNA or RNA molecule passes through a nanopore, the disruption in ionic current is measured, producing an electrical signal that corresponds to the specific nucleotides in the sequence [5]. These raw signals, representing complex biochemical interactions, are then processed by sophisticated algorithms and neural networks trained to recognize patterns in the signal data [6]. Neural networks excel at decoding these intricate patterns, converting the continuous stream of electrical signals into accurate nucleotide sequences. This integration of biophysical measurement with cutting-edge artificial intelligence enables rapid, real-time interpretation of genetic information, transforming raw data into meaningful genomic insights [7,8]. The concept of nanopore sequencing was first proposed in the mid-1990s, based on the idea of threading single DNA or RNA molecules through a tiny pore and detecting changes in ionic current to read the sequence [9]. Early research focused on biological nanopores, such as alpha-hemolysin, which provided proof of principle but faced significant technical challenges. It wasn’t until the late 2000s that significant advancements were made, particularly by Oxford Nanopore Technologies, which developed robust nanopore sequencing platforms [10]. Subsequent developments have improved the accuracy, throughput, and range of applications of nanopore sequencing, solidifying its role as a powerful tool in genomics research and diagnostics [11–13].

**Figure 1:**
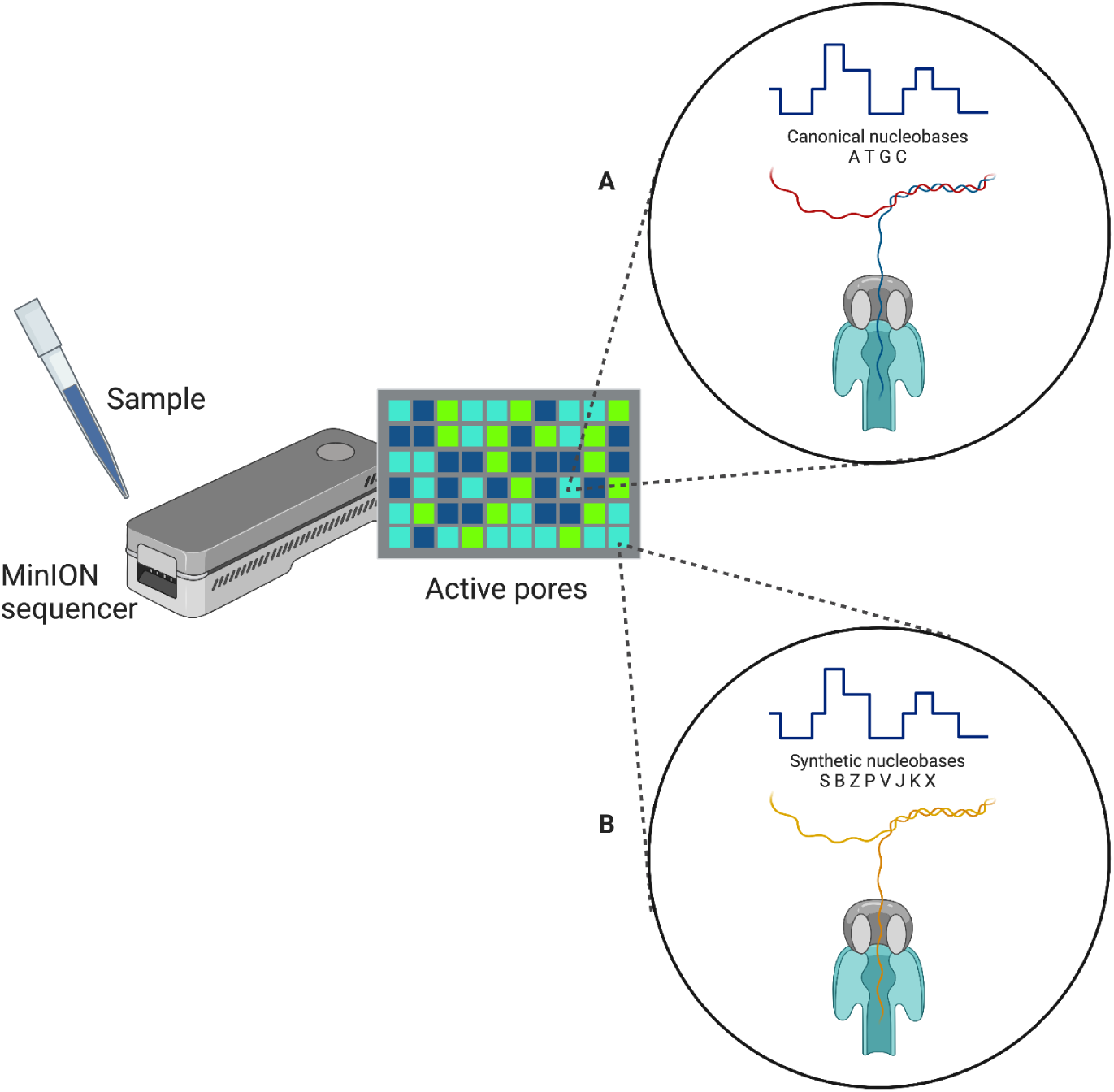
Principles of nanopore sequencing. (A) Nanopore sequencing of natural nucleobases found in DNA (Adenine, Thymine, Guanine, and Cytosine). (B) Nanopore sequencing of synthetic nucleobases such as the Artificially Expanded Genetic Information System (AEGIS). AEGIS is an engineered DNA analog system that uses twelve different nucleobases in its genetic code, including the four natural nucleobases plus eight synthetic nucleobases (S, B, Z, P, V, J, K, and X) that form S:B, Z:P, V:J and K:X base pairs.

In this study, we investigate the capacity of nanopore sequencing to decode the genetic information of parasitic organisms, such as protozoa and bacteriophages (phages), which possess hypervariable and dynamic genomes to evade host immunity. We utilize metagenomic samples from trypanosome-infected mice to test the limits of nanopore technology in directly decoding biological information from distinctive genomes within complex samples.

Trypanosomes are a group of protozoan parasites belonging to the genus *Trypanosoma*, which are responsible for several significant diseases in humans and animals [14,15]. African trypanosomiasis, commonly known as sleeping sickness, is caused by *Trypanosoma brucei* and transmitted by the tsetse fly and affects humans in sub-Saharan Africa [14,15]. Another notable disease is Chagas disease, caused by *Trypanosoma cruzi* and transmitted by triatomine bugs, primarily affecting people in Latin America [16]. These diseases are characterized by severe and often chronic symptoms, including fever, swelling, and neurological disorders, which can lead to death if left untreated [14,15]. Trypanosome genomes are difficult to study due to their complex structure, including large numbers of repetitive sequences and extensive gene duplication, which complicate genome assembly and annotation [17]. Additionally, they exhibit antigenic variation, where they periodically change their surface glycoproteins to evade the host’s immune system, adding another layer of complexity [14,15]. The presence of large subtelomeric regions prone to high rates of recombination further complicates genomic studies [18,19].

In this study, we demonstrate the ability of nanopore sequencing to detect and assemble the distinctive genomes of unicellular protozoa, even amidst the presence of other organisms such as phages, bacteria, and archaea in mice blood and fecal samples. This preliminary study showcases the remarkable potential of nanopore sequencing in the challenging task of classifying and assembling the *T. brucei* genome from metagenomic samples. Unlike short-read sequencing, nanopore technology generates long reads that can span repetitive and hypervariable regions, which are prevalent in the *T. brucei* genome. This capability enables more accurate and contiguous assemblies, overcoming the fragmentation issues associated with short-read methods [20–22]. The ability to classify and assemble such a complex genome from metagenomic samples underscores the robustness and versatility of nanopore sequencing, paving the way for more comprehensive studies on the genomic diversity and evolution of *T. brucei* and other similarly complex parasites [20–22]. By providing insights into distinctive biological entities, this study explores this novel technology as a future diagnostic tool for converting any type of biological information encoded on a sequence of organic materials into human-readable signals.

## Materials and Methods

### Sample preparation for nanopore sequencing: nucleic acid extraction and ligation

The experimental process of nanopore sequencing begins with the crucial step of nucleic acid extraction, which involves isolating high-quality DNA or RNA from the biological sample (Table S1). This step is critical because the integrity and purity of the nucleic acids directly affect the quality and quantity of the sequence output. Various methods can be used for extraction, depending on the source material and the downstream requirements [23]. Each method aims to remove contaminants such as proteins, lipids, and other cellular debris while ensuring that the nucleic acids are intact and free from inhibitors that could interfere with the sequencing process. High molecular weight DNA is particularly desirable for nanopore sequencing to maximize the read length capabilities of the technology [7,24].

Following nucleic acid extraction, the next key experimental element is the ligation of sequencing adapters to the extracted DNA or RNA [25]. This step involves attaching specific sequences, known as adapters, to the ends of the nucleic acid fragments. These adapters facilitate the interaction between the nucleic acids and the nanopore during sequencing. The ligation process typically uses a ligase enzyme to covalently bond the adapters to the nucleic acid ends. Successful adapter ligation is essential for efficient nanopore loading and translocation. In addition to standard adapters, barcoding adapters can be used to multiplex several samples in a single sequencing run, allowing for cost-effective and high-throughput sequencing [26]. After ligation, the prepared nucleic acid library is loaded onto the nanopore flow cell, where the sequencing occurs [27]. The accuracy and efficiency of the ligation step are critical, as any failures can lead to incomplete sequencing data or reduced sequencing quality.

In this study, we conducted infection and control experiments to collect blood and fecal samples from trypanosome-infected mice and control mice, respectively (Table S2). The infection experiments were conducted by infecting two mice with *Trypanosoma brucei* and collecting the fecal samples on the third day of infection and the blood samples on the fifth day of infection, which is regarded as the infection peak [28]. The control experiments were conducted by collecting the fecal samples on the third day and the blood samples on the fifth day from the healthy mice. Throughout the study, the samples from the mice blood are labeled as TrypOnly, TrypBlood_1, TrypBlood_2, and TrypBlood_Control, and the samples from the mice feces are labeled as Feces_Control and Feces_Infected. The blood sample with the label TrypOnly was processed to isolate the trypanosome parasites from the mice blood, and collected in the DPBS salt solution (Dulbecco’s Phosphate-Buffered Saline) at the concentration of 10^8^ cells in 500 µl.

From these experiments, we used a high-molecular-weight (HMW) DNA kit to extract DNAs from the metagenomic samples. The MagAttract HMW DNA Kit (QIAGEN N.V., Germany) is designed to facilitate the purification of high-molecular-weight DNA, specifically ranging from 100 to 200 kilobases (kb). This kit utilizes magnetic beads that selectively bind to DNA molecules, allowing for the removal of contaminants and inhibitors through a series of wash steps. For the blood samples, we used a blood-specific protocol for the blood samples of 200 μl. For the fecal samples, we adapted a tissue-specific protocol by mixing 25 mg feces into 200 μl PBS solution (Sigma-Aldrich, USA). Subsequently, the extracted DNA was ligated using the SQK-LSK110 specific for the MinIoN flowcells (FLO-MIN106) that require 1 µg (or 100-200 fmol) gDNA. We used a DNA Spectrophotometer (DeNovix Inc., USA) to check the quantity and quality of each extracted DNA (Table S1).

### Processing of nanopore sequence output: Quality control and basecalling

Downstream analysis of nanopore sequencing begins with quality control (QC) measures to ensure the integrity and reliability of the raw sequencing data [29]. QC involves evaluating various metrics such as read length distributions, read quality scores (Q-score), and yields of sequenced bases. Software tools like MinKNOW [30] and NanoPlot [31] provide visual and statistical summaries of these metrics, allowing researchers to identify and filter out low-quality reads or sequences that do not meet the predefined thresholds. Additionally, alignment tools such as Minimap2 [32] can be used to map the reads to a reference genome, helping to assess the overall accuracy and completeness of the sequencing run. This step is crucial for identifying potential issues related to the sequencing process, such as contamination or degradation of nucleic acids, which could compromise the downstream analyses.

Following QC, the next critical step in the downstream analysis is basecalling, which translates the raw electrical signals generated by the nanopore into nucleotide sequences. This process involves neural networks trained to recognize patterns in the ionic current disruptions caused by the passage of nucleotides through the nanopore [33]. The machine learning techniques adapted to the basecalling process include recurrent neural networks (RNN) such as DeepNano [34], convolutional neural networks (CNN) such as Causalcall [35] and Chiron [36], and transformer models such as Bonito [37]. There are several ways to estimate the Q-score of a reconstructed nucleotide from the patterns in the ionic currents generated during nanopore sequencing runs. For example, the Chiron and Causalcall tools calculate the Q-score using the ratio of the probability of occurrence of the most likely nucleotide (*p*1) over the probability of occurrence of the second likely nucleotide (*p*2) as follows: *Q_score_* = 10 · *log*_10_(*p*1/*p*2). On the other hand, the Bonito tool derives Q-scores directly from the trained model, using the probability of occurrence of the most likely nucleotide (*p*) and the internal variables (*x* and *y*) for scaling as follows: *Q_score_* = − 10 · (*y log*_10_ (1 − *p*). + *log*_10_ (*x*)) [33].

Basecalling software processes these signals in real-time or post-run, producing FASTQ files that contain the sequenced reads along with their Q-scores. Advances in machine learning have significantly improved basecalling accuracy, enabling the detection of not only the canonical nucleotides but also modified bases, which are essential for understanding epigenetic modifications [38]. The high-resolution data obtained through nanopore sequencing, coupled with robust basecalling, provides comprehensive insights into the genomic and epigenomic landscapes of the studied organisms [39]. In this study, MinION flowcells (FLO-MIN106) were run for at least 72 hours to maximize the sequence output and adaptive sampling was set to deplete the host genome of *Mus musculus* to enrich non-host genomes. We used the Guppy basecaller for real-time basecalling with the high-accuracy model and the Dorado basecaller for basecalling with the super-accuracy model after the experiments (Table S2). For these tools, each read can be filtered by a predetermined Q-score threshold set by the user, and we chose the minimum Q-score of 10 to filter out low-quality reads.

### Survey of nanopore sequence output: Taxonomic classification

Taxonomic classification of nanopore reads aims to identify and characterize the organisms present in a metagenomic sample [40,41]. This process involves assigning taxonomic labels to DNA sequences based on similarity-based approaches to reference databases or summary-based approaches that capture the unique sequence features of different taxa [42]. Tools like Kraken2 [43], Centrifuge [44], and MetaPhlAn [45] are commonly employed for taxonomic classification, each using different algorithms to match reads to known taxonomic groups. Kraken2, for instance, uses exact *k*-mer matches to a database of known sequences [43] and Centrifuge extends this by incorporating a compressed index of the entire NCBI nucleotide database, allowing for efficient and scalable taxonomic classification [44]. MetaPhlAn, on the other hand, focuses on marker genes specific to different taxa, offering high precision even with less comprehensive databases [45].

The accuracy of taxonomic classification is influenced by factors such as read length, sequencing accuracy, and the comprehensiveness of the reference database [42]. Nanopore reads, while typically longer than those generated by short-read technologies, have higher error rates, which can complicate the classification process [25]. To mitigate this, pre-processing steps such as error correction and filtering low-quality reads are often necessary [42].

Additionally, the use of databases that include a broad range of microbial genomes, including those from underrepresented or novel taxa, enhances the ability to classify reads accurately [7]. Once classified, the resulting taxonomic profiles can be analyzed to assess the diversity, composition, and potential functions of the microbial community [40,46]. This information is vital for understanding ecological relationships, disease mechanisms, and environmental impacts, thereby providing valuable insights into the biological and ecological significance of the studied samples [41,47,48].

For metagenomic identification, a single read can be used by searching against a specific database. For instance, the WIMP pipeline in the cloud-based EPI2ME platform enables rapid prokaryotic species identification from metagenomic samples using the Centrifuge classification [44] and the NCBI RefSeq database for bacteria, fungi, archaea, and viruses [49]. For reads above the quality threshold, the Centrifuge classification assigns each read to a taxon in the NCBI taxonomy. The WIMP reports counts of reads at given taxon ranks such as species, genus, and family, and generates a subtree representing the NCBI taxonomy. The Centrifuge engine classifies the read using mappings with at least one 22 bases (b) match and scores each species using an empirical formula that assigns greater weight to the longer segments:

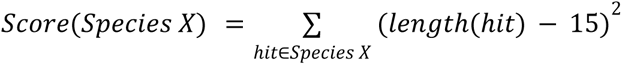

The formula considers that almost all sequences of 15 b or shorter match in the database by chance. For multiple species matches, the algorithm can be set to traverse up the taxonomic tree to output the lowest common ancestor of all matching species. Based on the comparison study, it is recommended to filter out the reads that score ≤300 and have match lengths ≤50 b to reduce false-positive assignments for error-prone reads [44].

We ran the metagenomic workflow using the classification method of Kraken2 in EPI2ME with a minimum read length of 200 bases. The EPI2ME has a collection of bioinformatics workflows that allow for the taxonomic classification of shotgun metagenomic data. It includes software such as Kraken2 [43] and Minimap2 [32] for the taxonomic classification of nanopore reads. Kraken utilizes a database of a *k*-mer and the last common ancestor (LCA) of all organisms whose genomes contain that *k*-mer to allow a quick search of the lowest LCA in the taxonomic tree associated with a target *k*-mer [50]. Minimap2 maps DNA or long mRNA sequences against a reference database for accurate short reads of ≥100 b in length or genomic reads of ≥1000 b in length at an error rate of ∼15% [32]. For the Kraken classification, we used the PlusPFP-8 [43] as the reference database, which includes archaea, bacteria, virus, plasmid, vector, protozoa, fungi, plant, and human, to include the broadest range of genomes available in the tool. To measure the alpha diversity of species in the samples, we used the Shannon index in the EPI2Me pipeline. After categorizing the resulting sequences into taxonomic bins, the number of species in each sample was calculated to determine the species richness. Next, the distribution of abundance across these species was calculated to assess whether the species were evenly distributed or if certain species were dominant.

### Analysis of nanopore sequence output: Metagenomic de novo assembly

Metagenomic assembly of nanopore reads involves reconstructing genomes from the mixed DNA sequences in environmental or clinical samples, bypassing the need for culturing individual species [51]. Metagenomic assembly using nanopore sequencing can capture extensive genomic regions, including repetitive elements and structural variations, that are often missed by short-read technologies [52–54]. Assembly tools such as Canu [55], Flye [56], and metaSPAdes [57] are commonly used to piece together the long reads into contiguous sequences (contigs). These tools must address challenges such as high error rates inherent to the earlier version of nanopore flow cells and the presence of highly diverse microbial populations with varying abundances. Strategies like read correction [58] and hybrid assembly (combining nanopore long reads with high-accuracy short reads) [59] were often employed in the earlier works to improve the accuracy and completeness of the assembled genomes.

Once the initial assembly is completed, additional steps are taken to refine and validate the metagenomic assemblies. Contigs are binned into individual genomes using binning algorithms like MetaBAT [60], MaxBin [61], or CONCOCT [62], which group sequences based on nucleotide composition and coverage patterns. These bins represent the draft genomes of different species present in the sample. Annotation tools such as Prokka [63] or RAST [64] can then be used to predict gene functions and identify metabolic pathways, providing insights into the ecological roles of the community members. Quality assessment of the assemblies involves metrics like N50, completeness, and contamination rates, typically evaluated using tools like CheckM [65] or QUAST [66]. The high-throughput nature of nanopore sequencing, coupled with sophisticated assembly and binning techniques, allows researchers to uncover the genetic and functional diversity of microbial communities, shedding light on their structure, dynamics, and potential interactions within their environments.

In this study, we attempted metagenomic de novo assembly of protozoa and phages using the advantages of long reads. Due to the long read length, the chance of obtaining longer contigs or full genomes from shotgun metagenomic data is not negligible, making it particularly advantageous for reference-free de novo assembly [22]. In this study, Flye was used to assemble long reads into contigs. These contigs were mapped against the reference genomes of *T. brucei* using a mapping software called MUMmer [67] as the target genome in these trypanosome-infected samples was known.

## Results

### Nanopore sequence output of high-molecular-weight DNA

We explored the potential of nanopore sequencers in sequencing high-molecular-weight DNAs from diverse metagenomic samples. We first verified the quality and quantity of the extracted DNA and the sequenced output from each sample was satisfactory. The HMW DNA extracted from the metagenomic samples was checked with the DNA spectrophotometer, which showed that while the concentrations were satisfactory, the quality metrics of 260/280 and 260/230 fell slightly outside the acceptable ranges of ∼1.8 and 2.0 to 2.2, respectively (Table S1). The 260/230 ratios were consistently low in the DNA extracted from the blood samples, indicating that they had contaminants absorbing at 230 nm or less. This contamination may result from the extraction technique of the MagAttract HMW DNA kit, which uses a magnetic bead-based method for removing contaminants and inhibitors. While this gentle purification is ideal for preserving the integrity of HMW DNAs, it may not be as effective at removing contaminants and inhibitors for certain sample types. Despite the suboptimal quality of the extracted DNAs, the subsequent nanopore sequencing runs yielded satisfactory sequence outputs in terms of both quality and quantity, as described below.

The read N50 value represents the mean length of a set of reads in nanopore sequencing, and the N50 values from this study show a wide range from 33,550 b to 582 b (Table 1). The blood samples tend to have shorter N50 values, suggesting the blood-specific DNA extraction protocol may adversely affect the length of the extracted DNA (Figure S1).

**Table 1:**
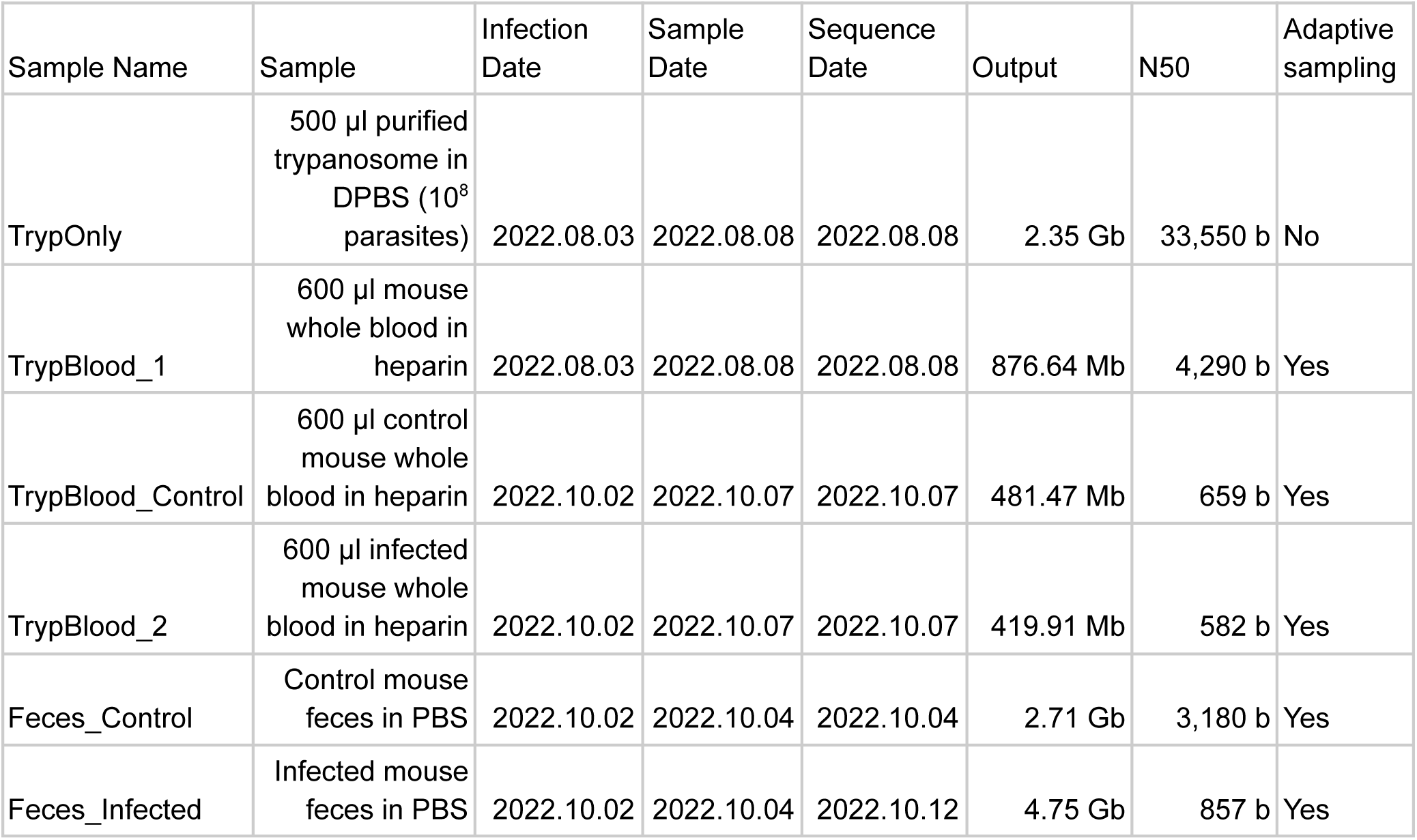
Experimental design and output of nanopore sequencing from the metagenomic samples containing host-dependent protozoa and phages. Adaptive sampling was done against the host reference genome of *mus musculus*.

| Sample Name | Sample | Infection Date | Sample Date | Sequence Date | Output | N50 | Adaptive sampling |
| --- | --- | --- | --- | --- | --- | --- | --- |
| TrypOnly | 500 µl purified trypanosome in DPBS (10<sup>8</sup> parasites) | 2022.08.03 | 2022.08.08 | 2022.08.08 | 2.35 Gb | 33,550 b | No |
| TrypBlood_1 | 600 µl mouse whole blood in heparin | 2022.08.03 | 2022.08.08 | 2022.08.08 | 876.64 Mb | 4,290 b | Yes |
| TrypBlood_Control | 600 µl control mouse whole blood in heparin | 2022.10.02 | 2022.10.07 | 2022.10.07 | 481.47 Mb | 659 b | Yes |
| TrypBlood_2 | 600 µl infected mouse whole blood in heparin | 2022.10.02 | 2022.10.07 | 2022.10.07 | 419.91 Mb | 582 b | Yes |
| Feces_Control | Control mouse feces in PBS | 2022.10.02 | 2022.10.04 | 2022.10.04 | 2.71 Gb | 3,180 b | Yes |
| Feces_Infected | Infected mouse feces in PBS | 2022.10.02 | 2022.10.04 | 2022.10.12 | 4.75 Gb | 857 b | Yes |

However, the purified trypanosome blood sample had the highest N50 value and the infected fecal sample also exhibited a relatively short N50 at 857 b, suggesting that these differences are more likely to be inherent to the genomic composition within each sample. The N50 values in some experiments were shorter than anticipated, despite utilizing the HMW DNA kit designed to purify genomes ranging from 100 to 200 kb. However, these values were still longer compared to those obtained using microbiome-specific DNA extraction kits with a mechanical lysis step [7].

The sequence output varied significantly across different sample types, with the blood samples consistently yielding lower output values below 1 gigabases (Gb) (Figure S2). This result supports the earlier hypothesis that the intrinsic composition of the blood samples differs from that of the other sample types. As expected, the fecal samples produced the most sequence output, with the control feces and the infected feces producing 2.71 Gb and 4.75 Gb, respectively (Table 1). The highest output produced from these nanopore sequencing runs reached 10% of the theoretical maximum output of a MinION flow cell (Table S2), a result consistent with observations from other studies [7,68].

### Taxonomic classification of nanopore reads

For basecalling, we utilized the advanced basecaller Dorado, employing the super-accuracy model to ensure the highest level of accuracy in our results [33]. The median Q-scores from all sequence outputs achieved above 10, which indicates the basecall accuracy of at least 90% (Figure S3). For reads with a Q-score above 10, we processed them through the metagenomic workflow using the Kraken classification method in EPI2ME, setting the minimum read length to 200 bases, based on the study showing the minimum read length of 200 bases ensures higher confidence in the classification [69]. Overall, the fecal samples contain more diverse organisms when compared to the blood samples (Figure 2). When the number of classified species in the superkingdoms of Archaea, Viruses, Eukaryota, and Bacteria was summarized in each sample, the fecal samples contained more than 10 folds of different species as compared to the blood samples (Table 2). As expected, this increased number of species is most notable in the superkingdom of Bacteria for both fecal samples (Feces_Control and Feces_Infected). For the blood samples, the control sample (TrypBlood_Control) had a lower number of species than the infected samples (TrypOnly, TrypBlood_1 and TrypBlood_2).

**Figure 2:**
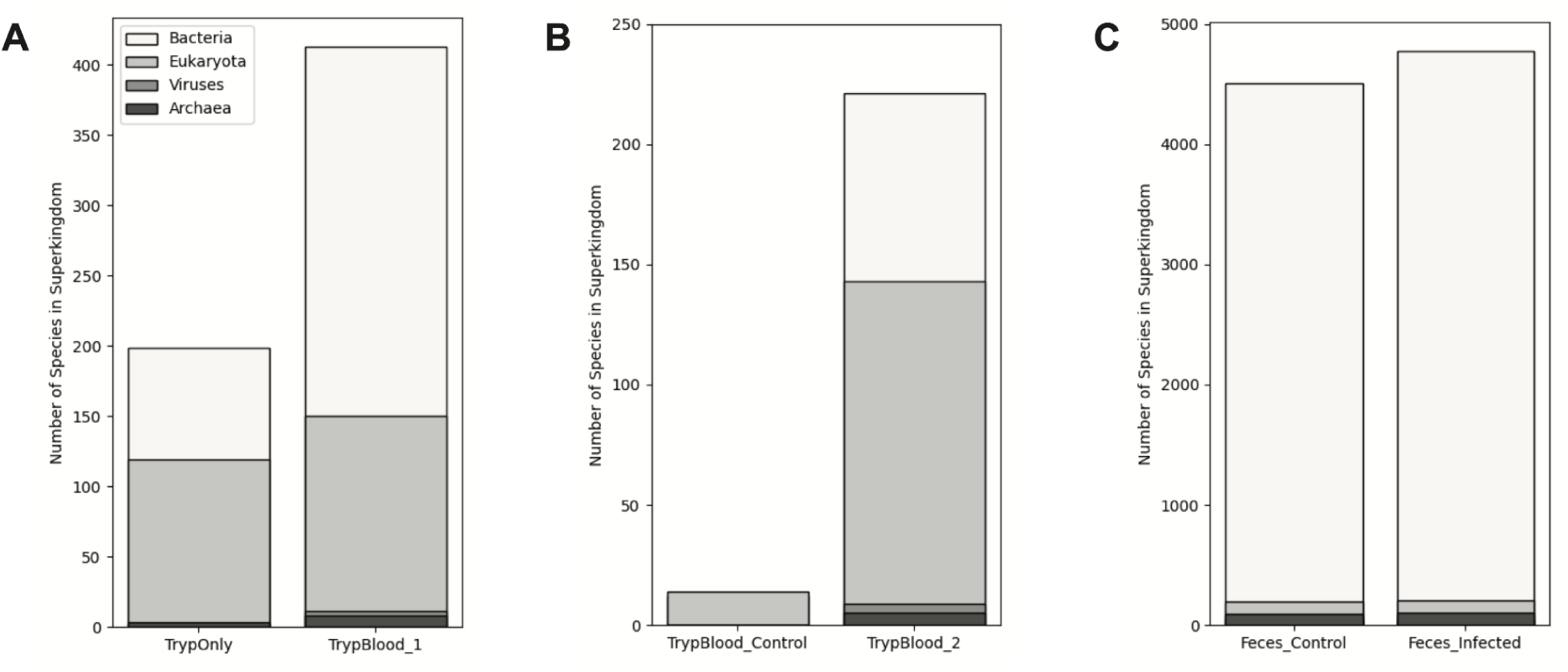
Comparison of the number of classified species in each superkingdom (Archaea, Viruses, Eukaryota, and Bacteria) between the control and infection experiments. (A) Number of classified species in each superkingdom for the trypanosome only sample (TrypOnly) and the trypanosome in blood sample (TrypBlood_1). (B) Number of classified species in each superkingdom for the trypanosome control sample (Tryp_Control) and the trypanosome in blood sample (TrypBlood_2). (C) Number of classified species in each superkingdom for the feces control sample (Feces_Control) and the feces infected sample (Feces_Infected).

**Table 2:**
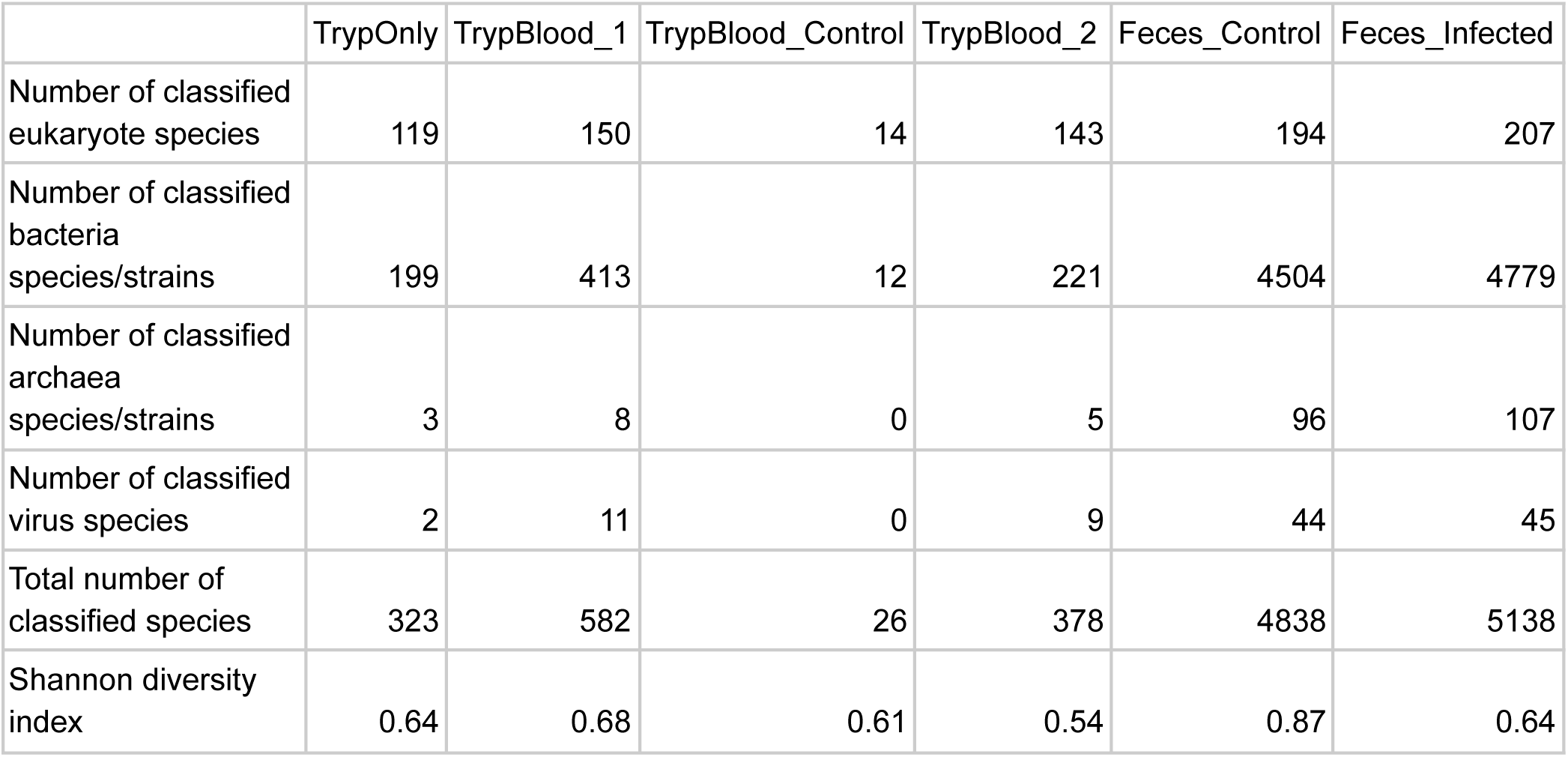
Taxonomical identification for each experiment using the Kraken classification method. The Shannon diversity index assesses the alpha diversity of bacterial samples. Higher values indicate a greater presence of rare species within the sample.

|  | TrypOnly | TrypBlood_1 | TrypBlood_Control | TrypBlood_2 | Feces_Control | Feces_Infected |
| --- | --- | --- | --- | --- | --- | --- |
| Number of classified eukaryote species | 119 | 150 | 14 | 143 | 194 | 207 |
| Number of classified bacteria species/strains | 199 | 413 | 12 | 221 | 4504 | 4779 |
| Number of classified archaea species/strains | 3 | 8 | 0 | 5 | 96 | 107 |
| Number of classified virus species | 2 | 11 | 0 | 9 | 44 | 45 |
| Total number of classified species | 323 | 582 | 26 | 378 | 4838 | 5138 |
| Shannon diversity index | 0.64 | 0.68 | 0.61 | 0.54 | 0.87 | 0.64 |

The species rank from the metagenomic taxonomic classification reports shows the relative abundance of the 10 most abundant species in each sample (Figure S4). In the blood samples, the isolated trypanosome sample in DPBS (TrypOnly) shows the highest abundance of *Trypanosoma brucei* at over 80%, while the trypanosome samples in blood (TrypBlood_1 and TrypBlood_2) show the second most abundance at less than 10%. As expected, the control blood sample (TrypBlood_Control) did not have *T. brucei* in the top 10 species rank.

In the fecal samples, the 10 most abundant species all belonged to the superkingdom of bacteria, except for *Homo sapiens*. All the samples had *Homo sapiens* as the 10 most abundant species, suggesting the potential contamination during the experimental runs. However, this may be an artifact of the Kraken classification method using the PlusPFP-8 reference database, where the reference genome of *Homo sapiens* is included but not that of *Mus musculus*. The percentage of reads classified as *Homo sapiens* from the Kraken classification further supports this hypothesis, as the percentage of *Homo sapiens* reads varies from sample to sample, indicating this may be an intrinsic property of each sample (Table S3). For instance, the percentage of *Homo sapiens* reads in the TrypBlood_Control sample is as high as 30% of the total reads while that in the Feces_Infected sample is as low as 0.3% of the total reads. This result is also consistent with the fact that blood samples are likely to contain more host cells than fecal samples.

### Statistical analysis of nanopore reads

The alpha diversity measuring the within-sample diversity of these metagenomic samples was calculated and visualized by the EPI2ME pipeline (Figure S5). The Shannon diversity index is a widely used metric for assessing the alpha diversity of bacterial samples. Higher Shannon diversity values indicate a greater presence of rare species within a sample. Notably, The Feces_Control sample demonstrates the highest Shannon diversity index at 0.87, while the TrypBlood_Control sample has a low Shannon diversity index at 0.61 (Table 2).

We further compared the read count per classified species in the control and infected samples (Figure 3). The distribution of read counts per classified species is visualized as a boxplot, showing the read count of each species with the read count above 20 and the read length above 200 bases (Figure 3A). The boxplot shows that there are more species and with higher read counts in the infected samples (TrypBlood_2 and Feces_Infected) as compared to those in the control samples (TrypBlood_Control and Feces_Control). Notably, the blood control sample (TrypBlood_Control) only has one species of *Homo sapiens* with the read count above 20 and the read length above 200 bases, possibly a result of contamination or mis-classification as explained in the previous section. The infected fecal sample (Feces_Infected) has the most widely distributed number of species, showing it has the most variable numbers of read counts. This sample also has the highest read count of classified species from these datasets at 60,000 reads for *Duncaniella sp*. (Figures 3C). *Duncaniella* is a bacterial genus recently discovered to be dominant in the mouse gut microbiota [70]. Interestingly, *Duncaniella dubosii* was the second most abundant species after *Escherichia coli* in the control fecal sample of Feces_Control (Figure 3B) [71]. Among the 10 most abundant species in the fecal samples, several other species overlap between the infected sample and the control sample, including *Duncaniella sp.*, *Muribaculum intestinale*, *Bacteroides caecimuris*, and *Escherichia coli* which are commonly found in mouse feces [72]. Interestingly, a Gram-positive bacterium named *Curtobacterium flaccumfaciens* that causes diseases in plants was found with very high read counts at 8234 and 27265 in the Feces_Control and Feces_Infected, respectively [73].

**Figure 3:**
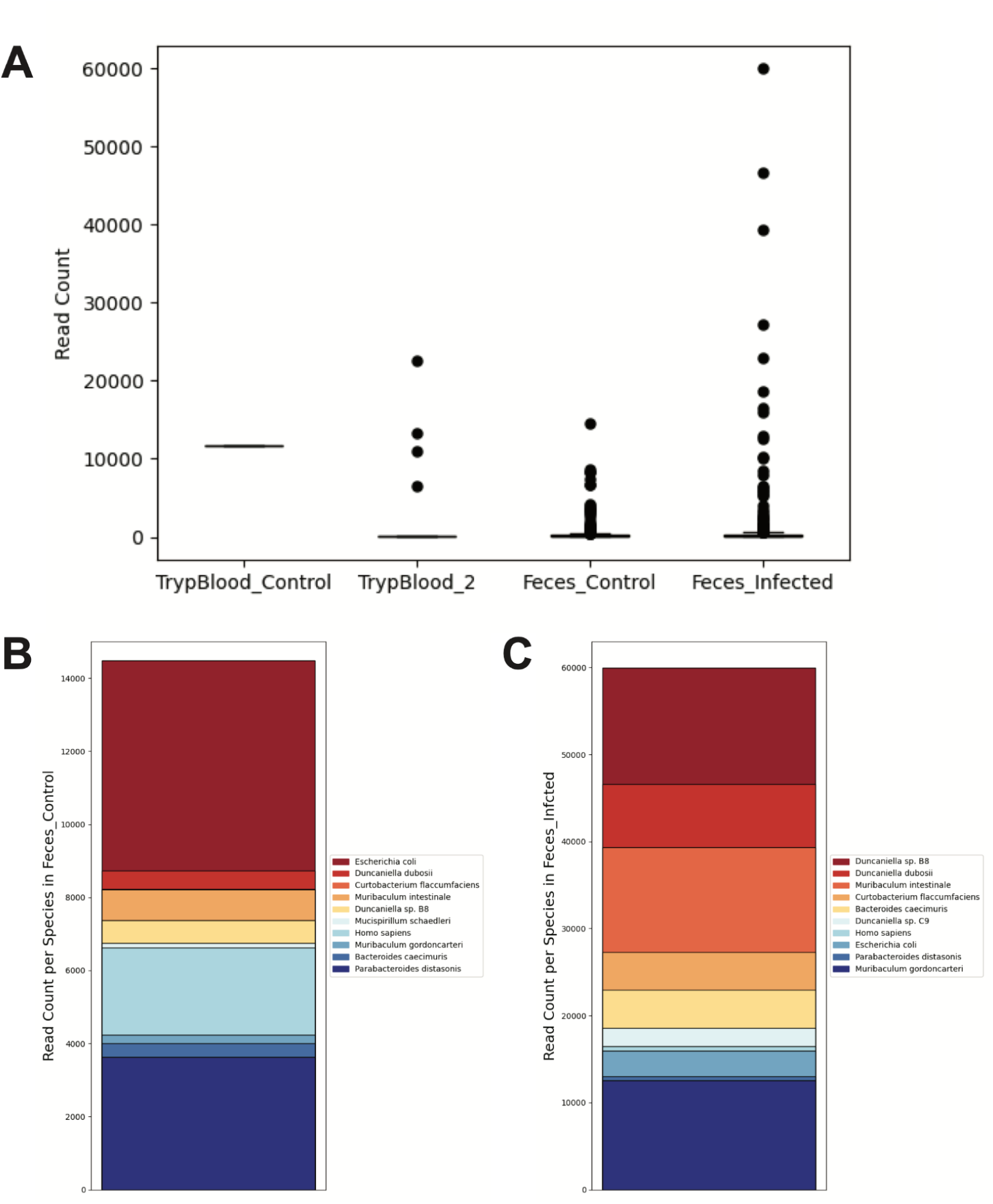
Comparison of read counts per classified species in the control samples and the infected samples. (A) The boxplot shows the distribution of read count per classified species above 20, with each dot representing the read count of each classified species, and the line showing the median of read counts of the classified species within the given sample. (B) The ten most abundant species rank in Feces_Control. (C) The ten most abundant species rank in Feces_Infected.

Next, we explored the relationship between the species count within the infected samples from the feces and blood collections (Figure 4). We included the species above the read count of 20 that were both identified in the blood and fecal samples (Figure 4A). A general linear regression shows that the species count in the infected fecal sample predicts the species count in the infected blood sample, with the R-squared value and *p*-value calculated as 0.89 and 1.69e^-10^ (Figure 4A). The relationship between the read count of each species present in both samples above the read count of 20 was positive, with the linear regression model given as *Blood_Infected* = 0. 37 * *Feces_Infected* − 3582. 26. Next, we investigated the low abundance region, where the read count of each species was below 1,000 (Figure 4B). When we focused on the low abundance area, a general linear regression still shows the species count in the control fecal sample predicts the species count in the control blood sample, with the R-squared value and *p*-value calculated as 0.52 and 3.29e^-61^ (Figure 4B). The relationship between the read count of each species present in both samples below the read count of 1,000 was positive, with the linear regression model given as

**Figure 4:**
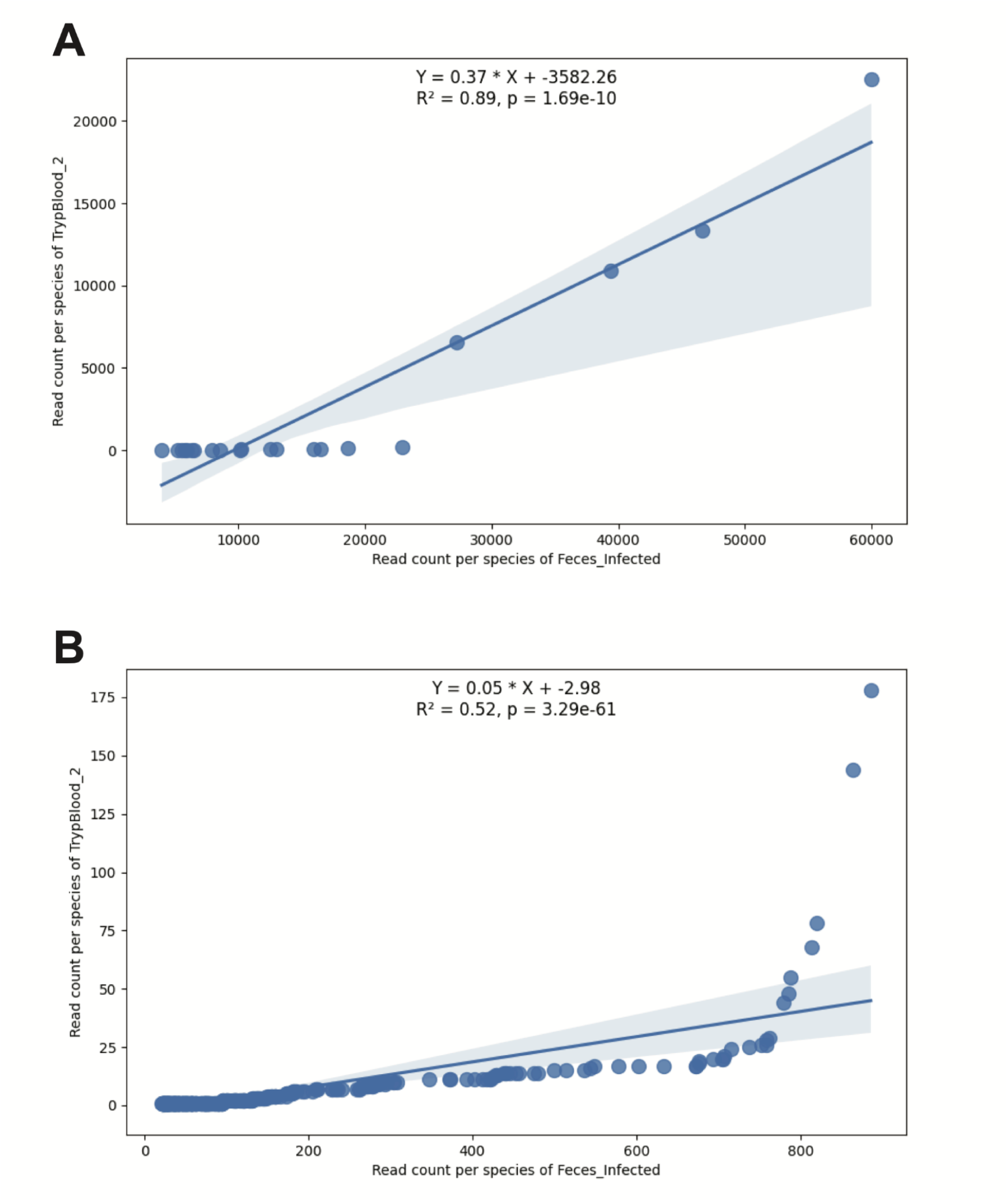
Relationship between the species count in the blood and fecal samples of the trypanosome-infected mice. Each data point is a species identified in the blood and fecal samples. A general linear regression was fitted to explore the relationship between the species count of the blood sample and that of the fecal sample. (A) Only species above the read count of 20 in the blood and fecal samples were included in the analysis. (B) Only species below the read count of 1000 in the blood and fecal samples were included in the analysis.

*Blood_Infected* = 0. 05 * *Feces_Infected* − 2. 98.

### Metagenomic de novo assembly of nanopore reads

We explored the capacity of nanopore sequencing in de novo assembly of distinctive genomes like *Trypanosoma brucei*, a protozoan parasite responsible for African sleeping sickness [14,15], from metagenomic samples. The *T. brucei genome* is highly complex and challenging to assemble using short-read sequencing technologies due to its extensive repetitive regions and intricate structure [18,19]. *T. brucei* has a diploid genome approximately 26 Mb in size, consisting of 11 pairs of megabase chromosomes, intermediate chromosomes, and a multitude of minichromosomes. The genome is characterized by a significant amount of repetitive DNA, including numerous transposable elements and large gene families, such as the Variant Surface Glycoprotein (VSG) genes, which are critical for immune evasion. These repetitive sequences create ambiguities in short-read sequencing, as short reads cannot adequately span and resolve these repetitive regions, leading to fragmented and incomplete assemblies [20].

Additionally, the dynamic nature of VSG gene recombination and the presence of subtelomeric regions, which are prone to rearrangements, further complicate the assembly process [18,19]. Another distinctive feature of the *T. brucei* genome is kinetoplastids, which are the mitochondria in trypanosomes and other related protozoa, containing their own DNA known as kinetoplast DNA (kDNA) [74]. They are highly structured network of circular DNA molecules found within the kinetoplast, consisting of thousands of interlocked minicircles (each 1-2 kilobases in size) and dozens of maxicircles (each 20-40 kilobases in size), playing a crucial role in mitochondrial function and the parasite’s life cycle.

The *T. brucei* genome is highly dynamic with frequent events, especially in the VSG gene arrays, and has abundant repetitive sequences, particularly in the minichromosomes and subtelomeric regions, making it an ideal benchmark to test the capacity of de novo assembly using long-read sequencing. Using the metagenomic option, the long reads of each sample were assembled de novo into the contigs (Table S4). The fecal samples showed high numbers of contigs, with the Feces_Control and Feces_Infected showing more than 10,991 and 8,903 assembled fragments, respectively. The blood samples all showed lower numbers of contigs, but the fragment N50 has higher lengths, particularly the TrypOnly sample with 68,574 bases in length.

With the hypothesis that some of these assembled fragments may be *T. brucei*, we mapped them against the reference genome of *T. brucei*, with the total genome length of 26.1 Mb that include 11 pairs of megabase chromosomes. The mapping results show that many contigs mapped to the reference genome of *T. brucei*, particularly in the infected blood samples (Figure 5). The infected blood samples of TrypOnly, TrypBlood_1, and TrypBlood2 had 98.61%, 97.08%, and 26.81% of the aligned bases to the reference genome of *T. brucei*, respectively (Table S5). These percentages were much lower in the other samples, where less than 0.03% of the reference genome of *T. brucei*, was aligned by the assembled fragments in TrypBlood_Conrol, Feces_Control, and Feces_Infected (Table S5). In terms of accuracy, the results showed high values of average identity in the infected samples at around 96%, whereas lower values at around 93% in the other samples.

**Figure 5:**
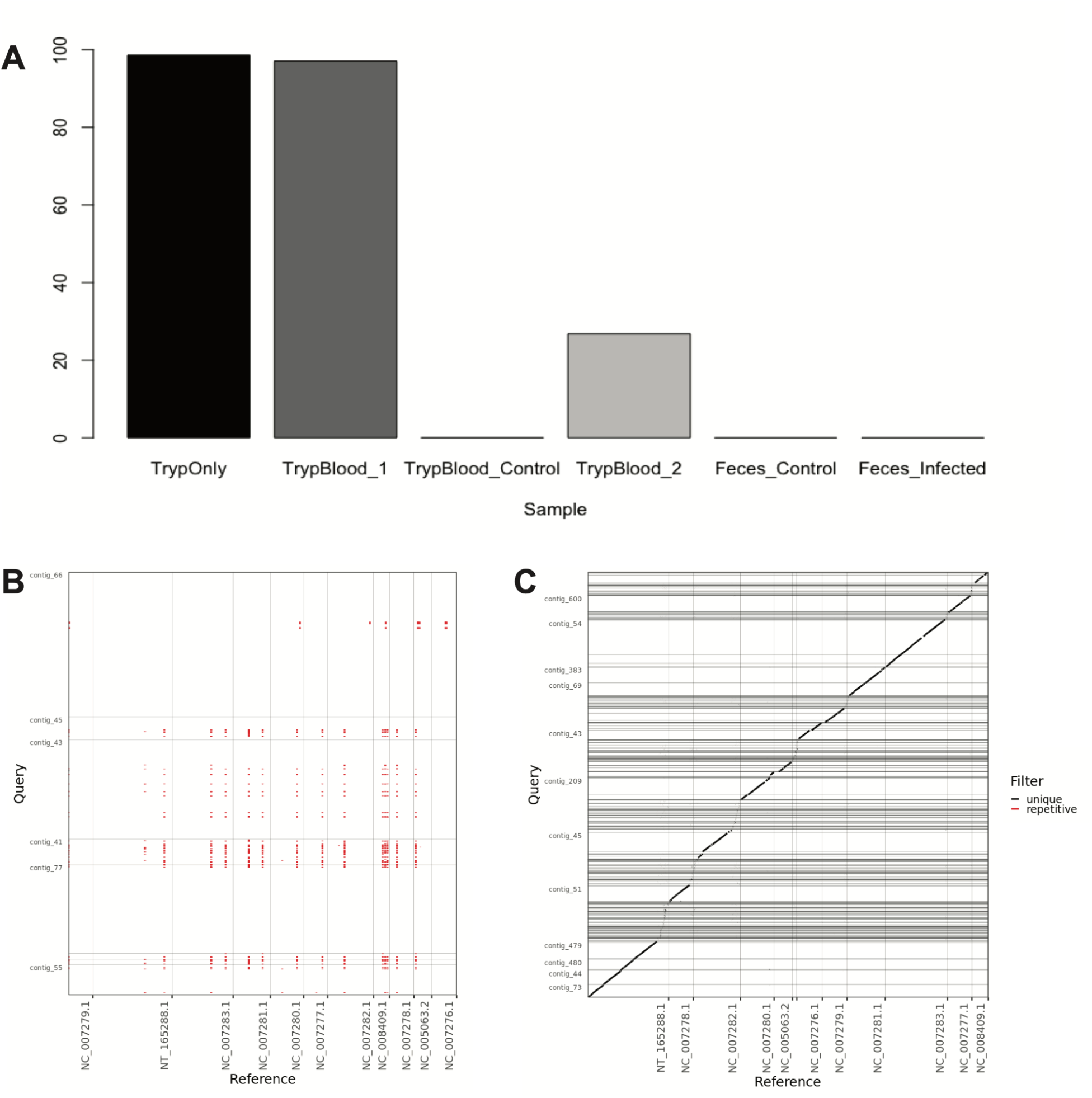
Trypanosome genome assembly visualization. (A) Percentage of the reference genome of *Trypanosoma brucei* aligned by the assembled fragments from each sample. (B) Dot plot of Assemblytics filtered alignments from the control experiment of TrypBlood_Control. (C) Dot plot of Assemblytics filtered alignments from the infected experiment of TrypBlood_1.

## Discussion

Nanopore sequencing technology, with its ability to read long sequences of nucleic acids by detecting changes in electric currents as molecules pass through nanopores, has the potential to decode a wide range of biologically encoded information [75]. Recently, nanopore sequencing has contributed to the expansion of human genomes [52,54] and the cell-free DNA profiling [76–78]. Nanopore sequencing has also been applied to various diagnostic settings to detect bacterial and viral pathogens, as well as drug resistance and vaccine target surveillance [79–82]. Thanks to the innovation of decoding biological information as an ionic current through a nanopore, this technology has the potential to extend beyond traditional nucleic acid and amino acid sequences. For instance, it may be adapted to read sequences of other biochemical molecules, such as those found in Artificially Expanded Genetic Information Systems (AEGIS) [83]. AEGIS expands the genetic alphabet by incorporating non-standard nucleotides, which can encode additional information beyond the canonical A-T and G-C base pairs. The sensitivity of nanopore sequencing to variations in electrical signals has the potential to distinguish these novel bases, providing a means to accurately decode these synthetic genetic systems (Figure 1B). This capability extends the utility of nanopore sequencing from biological molecules to engineered or extraterrestrial systems, enabling innovative applications in biotechnology, synthetic biology, and space biology, where information can be decoded from a broader array of molecular sequences [22].

Furthermore, the ability to directly sequence long strands of DNA or RNA without the need for fragmentation and amplification enables comprehensive insights into complex genomic regions, structural variants, and epigenetic modifications [21]. This direct sequencing approach significantly reduces errors associated with short-read sequencing and allows for a more accurate and contiguous assembly of genomes, making it an invaluable tool for studying distinctive biological systems [21]. In this study, we tested the capacity of nanopore sequencing in investigating hypervariable and dynamic genomes from metagenomic samples. We collected blood and feces samples during ongoing experimental studies of the trypanosome infection in mice. Trypanosome genomes are characterized by extensive variability and structural complexity, including numerous repetitive elements, gene duplications, and large-scale rearrangements [18,19,28]. Short-read sequencing, while highly accurate for contiguous regions, struggles to capture these complex genomic architectures due to its limited read lengths. Long-read sequencing technologies can produce reads that encompass these regions, which are necessary to achieve a more accurate and contiguous assembly of highly complex parasite genome [84].

In this study, we sequenced three sets of metagenomic samples using the MinION flowcells with the maximum capacity to generate 50 Gb of nanopore sequence output. The taxonomic classification of these nanopore sequence outputs identified a high proportion of reads as trypanosome genomes in the infected blood samples (Figure 5). The control blood samples and the fecal samples did not contain significant amounts of reads classified as trypanosome genomes, a result consistent with observations from other trypanosome infection studies [85]. The metagenomic de novo assembly of these nanopore sequence output from the infected blood samples shows that the reads identified as trypanosome genomes were successfully assembled into contigs that mapped on the reference genome of *Trypanosoma brucei* with an accuracy above 96%. Given the complex nature of trypanosome genomes, this accuracy represents a promising preliminary step for utilizing long-read sequencing in metagenomic de novo assembly of distinctive genomes such as *T. brucei*. However, more work is needed for finer resolution analysis, such as considering single-nucleotide polymorphisms (SNPs) or pangenome studies [86].

This study, while promising, has several limitations that must be addressed to draw robust biological insights. A key limitation is the lack of biological and technical replicates, which are essential for validating the observed differences in microbial compositions between control and infected samples, particularly in the mice feces (Figure 3). Without replicates, it is difficult to discern whether the observed differences are due to biological variations or experimental variations. Furthermore, the inherent variability in microbial communities necessitates multiple replicates to account for stochastic fluctuations and ensure statistical significance [87,88]. We explored the statistical relationship between the species count in the infected fecal sample and the infected blood sample for the species that are present in both types of samples (Figure 4). The statistical analysis shows that the species count in the blood and fecal samples of the trypanosome-infected mice has a statistically significant linear relationship with the *p*-value less than 0.001. However, the study’s reliance on a limited set of samples also restricts the generalizability of the findings. Future research should incorporate a well-designed set of biological and technical replicates to enhance the reliability and interpretability of the data, ultimately leading to more definitive insights into the microbial dynamics associated with the trypanosome infection. These directions will collectively advance our understanding of the biology of distinctive organisms in complex samples, paving the way for novel therapeutic strategies and diagnostic applications.

## Declarations

### Data Availability Statement

All codes related to this project are available under an open-source license at https://github.com/hshimlab. For data acquisition, we used MinION Mk1B and FLO-MIN106. For data processing, we used MinKNOW v.22.05.5 and EPI2ME-labs v2.9.2. For data analysis, R v.3.6.0, Python v.3.12.1, Matplotlib v.3.8.0, pandas v.2.2.0, and seaborn v.0.13.1 were used. For genome assembly, we used Flye v2.9.2, MUMmer v4.0+, and Assemblytics.

### Ethics Statement

The study was carried out in accordance with the recommendations of the ethical standards defined by the Declaration of Helsinki, and approved by the Institutional Review Board of GUGC (IACUC 2022-008).

### Author Contributions

Experiments were primarily conducted by H.S. Analyses were primarily conducted by B.H. and H.S. Specifically, nanopore analyses were performed by H.S., and genome analyses and visualization analyses were led by B.H. The study was conceived by H.S., and all authors contributed to writing the manuscript.

## Acknowledgments

The research and development activities described in this study were funded by GUGC and CSU, Fresno. The authors would like to thank S. Depuydt and M. Radwanska for providing the trypanosome samples and the members of the Centre for Biotech Data Science at GUGC. The authors would also like to thank X. Her and O. Emmanuel for inspiring the visualization and the members of the Department of Biology at California State University, Fresno.

